# Cooperative policing behavior regulates reproductive division of labor in a termite

**DOI:** 10.1101/2020.02.04.934315

**Authors:** Qian Sun, Jordan D. Hampton, Kenneth F. Haynes, Austin Merchant, Xuguo Zhou

**Affiliations:** Department of Entomology, Louisiana State University Agricultural Center, Baton Rouge, LA 70803, USA; Department of Entomology, University of Kentucky, Lexington, KY 40546-0091, USA

**Keywords:** policing behavior, reproductive conflict, neotenic reproduction, termites, eusociality

## Abstract

Reproductive conflicts are common in insect societies where helping castes retain reproductive potential. One of the mechanisms regulating the conflicts is policing, a coercive behavior that reduces direct reproduction by other individuals. In eusocial Hymenoptera (ants, bees, and wasps), workers or the queen act aggressively toward fertile workers, or destroy their eggs. In many termite species (order Blattodea), upon the death of primary queen and king, workers or nymphs can differentiate into neotenic reproductives and inherit the breeding position. During this process, competition among neotenics is inevitable, but how this conflict is resolved remains unclear. Here, we report a policing behavior that regulates reproductive division of labor in the eastern subterranean termite, *Reticulitermes flavipes*. Our results demonstrate that the policing behavior is a cooperative effort performed sequentially by successful neotenics and workers. A neotenic reproductive initiates the attack of the fellow neotenic by biting and displays alarm behavior. Workers are then recruited to cannibalize the injured neotenic. Furthermore, the initiation of policing is age-dependent, with older reproductives attacking younger ones, thereby inheriting the reproductive position. This study provides empirical evidence of policing behavior in termites, which represents a convergent trait shared between eusocial Hymenoptera and Blattodea.

## 1. Introduction

Eusocial insects exhibit reproductive division of labor between a few reproductive individuals and numerous sterile workers. However, potential conflicts arise in species where workers are capable of reproduction [1-4]. In addition to queen pheromones that suppress worker reproduction in many species [5-7], policing behavior is an important solution to maintain reproductive harmony [2, 8]. Queen policing, a term coined by Oster and Wilson in 1978, describes behaviors carried out by the queen to retain her reproductive dominance over workers [9]. Worker policing, which was named by Ratnieks in 1988, was used to describe the actions of workers that reduce worker-produced males in favor of sons of the queen in honey bee [8]. The concept of “policing behavior” was expanded by Monnin and Ratnieks in 2001 to include all “coercive actions that reduce direct reproduction by other individuals”, which accommodates various forms of behavioral regulation observed in diverse social insects [10]. In eusocial Hymenoptera (ants, bees, and wasps), policing behavior is performed via egg-eating or different forms of aggression, such as immobilization, biting, and stinging [10-13]. Policing interactions may occur among workers [11], among reproductives [14, 15], or between reproductives and workers [12, 16-18]. However, the vast majority of investigations on policing behavior have been focused on Hymenoptera, with little is known about its occurrence or nature in termites (order Blattodea), a group of eusocial insects evolved 50 million years earlier than eusocial Hymenoptera [19].

Policing behavior serves two functions that are not mutually exclusive, which are regulating genetic conflicts and improving colony efficiency [8, 20, 21]. Genetic conflicts arise between colony members due to relatedness asymmetries. For example, in social Hymenoptera with haplodiploid sex determination, workers are often capable of laying unfertilized male eggs, and they are more closely related to their own sons than males produced by other individuals [2, 8, 22]. Moreover, policing behavior contributes to colony efficiency even when little or no genetic conflict is present, as it optimizes the allocation of colony resources to reproduction [8, 23], or maintains an adaptive colony-level phenotype [24].

Unlike social Hymenoptera, termite colonies are usually founded and dominated by a pair of primary reproductives (queen and king). Upon their death, neotenic reproductives of both sexes can differentiate either from workers, which become ergatoid reproductives, or from nymphs, which become nymphoid reproductives [25]. Reproductive succession by neotenics has been reported in at least 13.4% of “higher” termite genera (Termitidae) and 61.7% of “lower” termite genera (all other termite families) [26]. In termites with diplodiploid sex-determination, individuals are more closely related to their own offspring than that of their siblings, regardless of outbreeding or inbreeding. Therefore, genetic conflicts are potential and competition for reproduction between nestmates is expected [27]. Moreover, given the fact that workers are reproductively totipotent in many species, colony-level efficiency can be compromised if excess neotenics differentiate, as it results in reduced labor force and increased resource demand by the reproductives and their brood.

In the presence of fertile reproductives, neotenic formation is inhibited through pheromones [6, 28, 29], and policing behavior through overt aggression was considered rare in termites [28, 30]. However, during the process of reproductive succession when colonies are orphaned and inhibitory pheromones are temporarily absent, production of excessive neotenic reproductives in the colony is expected. In addition, cannibalism of neotenics was observed in several termite families including Termopsidae [31], Kalotermitidae [30, 32-34] and Rhinotermitidae [35, 36], suggesting the presence of policing behavior that directly regulates reproduction in termites. Empirical studies of the process and causes of policing behavior, however, are lacking. In this study, we conducted a series of experiments to understand whether the number of ergatoid neotenics is regulated behaviorally during reproductive succession, and how policing behavior is performed in the eastern subterranean termite, *Reticulitermes flavipes*. Furthermore, we investigated a proximate factor that determines the succession of reproductives.

## 2. Methods

### (a) Insect collection and maintenance

Field and laboratory colonies of *R. flavipes* were used in this study. Field colonies were collected from Red River Gorge area, Daniel Boone National Forest (Slade, Kentucky, USA) and the University of Kentucky Arboretum (Lexington, Kentucky, USA). These colonies were obtained in summer using trapping stations filled with dampened cardboard rolls. Once captured termites were extracted from traps, placed in Petri dishes (14.5 cm × 2.0 cm) with moistened unbleached paper towel as their only food source for 7-10 days before they were used in experiments. Only workers and soldiers were collected from the field. Laboratory colonies were established in 2010 by pairs of sibling alates from a dispersal flight in Lexington, Kentucky, and they were kept in closed plastic boxes filled with moistened wood mulch and pinewood blocks in laboratory for 5 years before use. All colonies were maintained in complete darkness (L:D = 0:24), at 27 ± 1ºC, 80%-99% RH.

### (b) Orphaning assay to test ergatoid number restriction

This assay was used to simulate the reproductive replacement process after the death of primary reproductives. Groups of 100 workers were kept in petri dishes (35-mm-diameter) with moistened paper towel placed at the bottom. Two treatments including “removal” and “non-removal” of ergatoids were conducted. This experiment was specifically designed in a way to compare between the numbers of ergatoids that can potentially differentiate and survive (“removal”) and that actually survived (“non-removal”). All termites were maintained at 27 ± 1ºC and in complete darkness for 90 days. Each dish was checked by identifying the sex and counting the number of newly differentiated ergatoids. Each ergatoid was removed and replaced with a worker in the “removal” treatment, but returned to the dishes in the “non-removal” treatment. Dishes in “removal” treatment were checked every day, while dishes in “non-removal” treatment were checked every 10 days to reduce stress to reproductives resulting from manipulations needed for sex identification. The number of remaining termites in each group was counted every 30 days, and mortality was calculated based on the difference between the numbers of initial and remaining individuals. Injured ergatoids were not counted. A total of 20 replications were made with five replications in each of the four colonies. Two field and two laboratory colonies were used in this experiment.

### (c) Orphaning assay for observation of policing behavior

This assay was designed to observe policing behavior under orphaning condition, which resembled the “non-removal” treatment above. The dishes were incubated at 27 ± 1ºC for a total of 90 days. Once a week, the dishes were checked for dryness, and water was added if the paper towels at the bottom appeared to be dry. Between 60 and 90 days, each dish was checked for the presence of ergatoids, and the dishes with ergatoids were selected to be video recorded for 6 days. Video cameras (Canon Vixia HF G20, Canon Inc., Tokyo, Japan) were used for recording and yield high quality images. The dishes and the cameras were shaded under a piece of cardboard (1.5 m × 0.8 m) during recording. Ergatoids were identified as males or females and color marked using enamel paint (Testor Corporation, Rockford, IL, USA) on their head capsule prior to recording. During recording, the dishes were checked every day for missing ergatoids (which were cannibalized), and newly formed ergatoids were color marked. When a marked ergatoid was missing, the video of the previous 24 hours was quantitatively analyzed.

We define the “victim” as the ergatoid that was eventually cannibalized, the “attacker” as the individual who attacked the victim, and “bystander” as the ergatoids who did not perform the first major attack. A major attack was recognized when the attacker visibly injured the victim such that the researcher could see the abdomen was torn, hemolymph was leaking, and the victim quickly fled. The first major attack was designated as time “0”, and the frequency of vibration and number of workers surrounding the victim were documented. A one-minute sample (30 seconds before and after the time point) was analyzed for all these behaviors, with the sample selected for the time midpoints 5, 15, 20, 25, 30, 45, and 60 minutes before and after time “0”. Worker density near the victim was an indicator of cannibalism. The density was quantified by counting the number of workers and soldiers within 4 mm radius from the center of the victim (the radius approximately equals to body length of a worker). Only field-collected colonies were used in this experiment.

### (d) Policing assay to test the effect of ergatoid age

This assay was used to determine if policing behavior in *R. flavipes* is associated with the age of ergatoids. Each group of 50 workers was kept in a petri dish (55-mm-diameter) lined with 6 layers of moistened paper towel. A pair of virgin ergatoids, female and male, was added to each dish on day 1. Female and male treatments were conducted, with a younger female or male ergatoid being added to the initial group every day. When added to the dish, the initial pair of ergatoids was 7 days post differentiation (7-day old) and became older over the course of the experiment, and subsequent ergatoids were no more than 7-day old. All of the ergatoids were color marked as previously described, and no new ergatoids differentiated during this assay. The dishes were recorded until one of the ergatoids was missing. The video was then analyzed to identify who attacked that ergatoid. A total of 9 and 10 replications were made for female and male treatments, respectively; aggressively interacting pairs that were the same sex were analyzed for their age differences. This experiment did not attempt to address sex-specificity, but it was designed for increased chance of sex-specific aggression, and only same-sex aggressions were analyzed to eliminate confounding factors associated with sex. Only field-collected colonies were used in this experiment.

### (e) Statistical analyses

Data were analyzed using R (https://www.r-project.org/) and Statistix 10 (Analytical Software, Tallahassee, FL, USA), and graphs were generated using SigmaPlot 13 (Systat Software Inc, Chicago, IL, USA). Data for ergatoid number and mortality in reproductive “removal” and “non-removal” assays were fitted to Poisson family generalized linear mixed models using R’s *glmer* function. In the model for ergatoid number, ergatoid sex and treatment group (removal vs. non-removal) were coded as fixed effects, while colony of origin was coded as a random effect. An observation-level random effect was introduced to avoid overdispersion. In the model for mortality, treatment group was coded as a fixed effect, while colony of origin was coded as a random effect. In both cases, data were analyzed separately for each 10-day interval. Data testing the influence of ergatoid age on aggressive interactions in the policing assay were analyzed in Statistix using a Wilcoxon signed-rank test.

## 3. Results

### (a) The number of ergatoids is restricted

In “removal” treatment, there were significantly more female and male ergatoids differentiated than that remained in “non-removal” treatment within 90 days (Figure 1; x^2^ = 137.14, df = 1, *P* < 0.001; GLMM, Poisson family; n = 20). At the end of day 90, there were 11.80 ± 2.59 (mean ± SE) female and 4.75 ± 1.06 male ergatoids differentiated when they were removed daily, compared to only 2.60 ± 0.34 female and 1.05 ± 0.05 male ergatoids if they were not removed. In addition, there was a significantly higher overall mortality in “non-removal” than that in “removal” treatment (Figure S1; x^2^ = 30.36, df = 1, *P* < 0.001; GLMM, Poisson family; n = 20). In “non-removal” groups, we frequently observed injured ergatoids partially consumed by workers, along with other intact ergatoids. Between the two treatments, the mortality difference (8.6 individuals at day 60 and 10.2 at day 90) closely matched the difference in the number of ergatoids that differentiated and that survived (6.2 at day 60 and 12.9 at day 90), suggesting cannibalism of ergatoids in “non-removal” treatment is primarily responsible for the difference in mortality.

**Figure 1.**
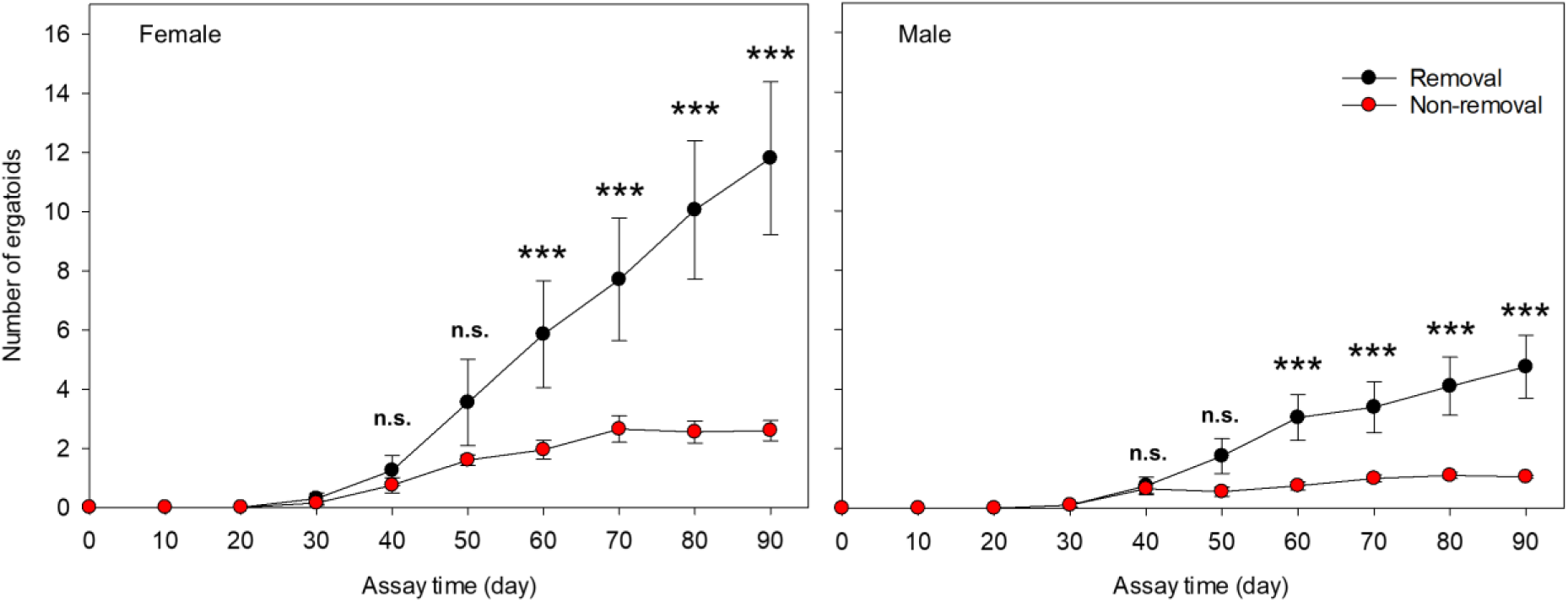
Restriction of ergatoid numbers in orphaning assay. Female and male ergatoids that formed from groups of 100 workers were documented for 90 days post establishment of experiment. Cumulative number of differentiated ergatoids (removal treatment) and surviving number of ergatoids (non-removal treatment) are presented (mean ± SE) and compared within the same observation day. n.s., not significant; ***, *P* < 0.001; GLMM, Poisson family; n = 20 per treatment per observation day.

### (b) Ergatoids and workers cooperate in elimination of excessive ergatoids

A total of seven events were captured with full behavioral process, which started with one ergatoid attacking another ergatoid and ended with the injured individual being cannibalized by workers (Figure 2, Table S1, Movies S1-S4). In this behavior, the attacker antennated the victim first, and used mouthparts to hold the abdomen or thorax of the victim before biting. The bite always caused the victim to leak hemolymph and quickly flee. Right after the attack, the attacker displayed alarm behavior by vigorously vibrating the body toward multiple directions; interestingly, the ergatoids who did not participate in the aggression (i.e., bystanders) also performed alarm behavior after the attack, while the victim rarely engaged in vibration (Figure 3a). Workers, on average, displayed little vibration (Figure 3a). With the alarm of ergatoids, workers rapidly began to surround the injured victim, biting and consuming it while it was still alive, and the cannibalism reached a peak 30 minutes after the attack (Figure 3b). As the cannibalism began to decline due to little of the remaining body to consume, the ergatoid attacker and bystanders began to decline in the frequency of alarm (45 to 60 minutes after the attack) (Figure 3a). Soldiers did not participate in either alarm behavior or cannibalism (Figure 3a, 3b).

**Figure 2.**
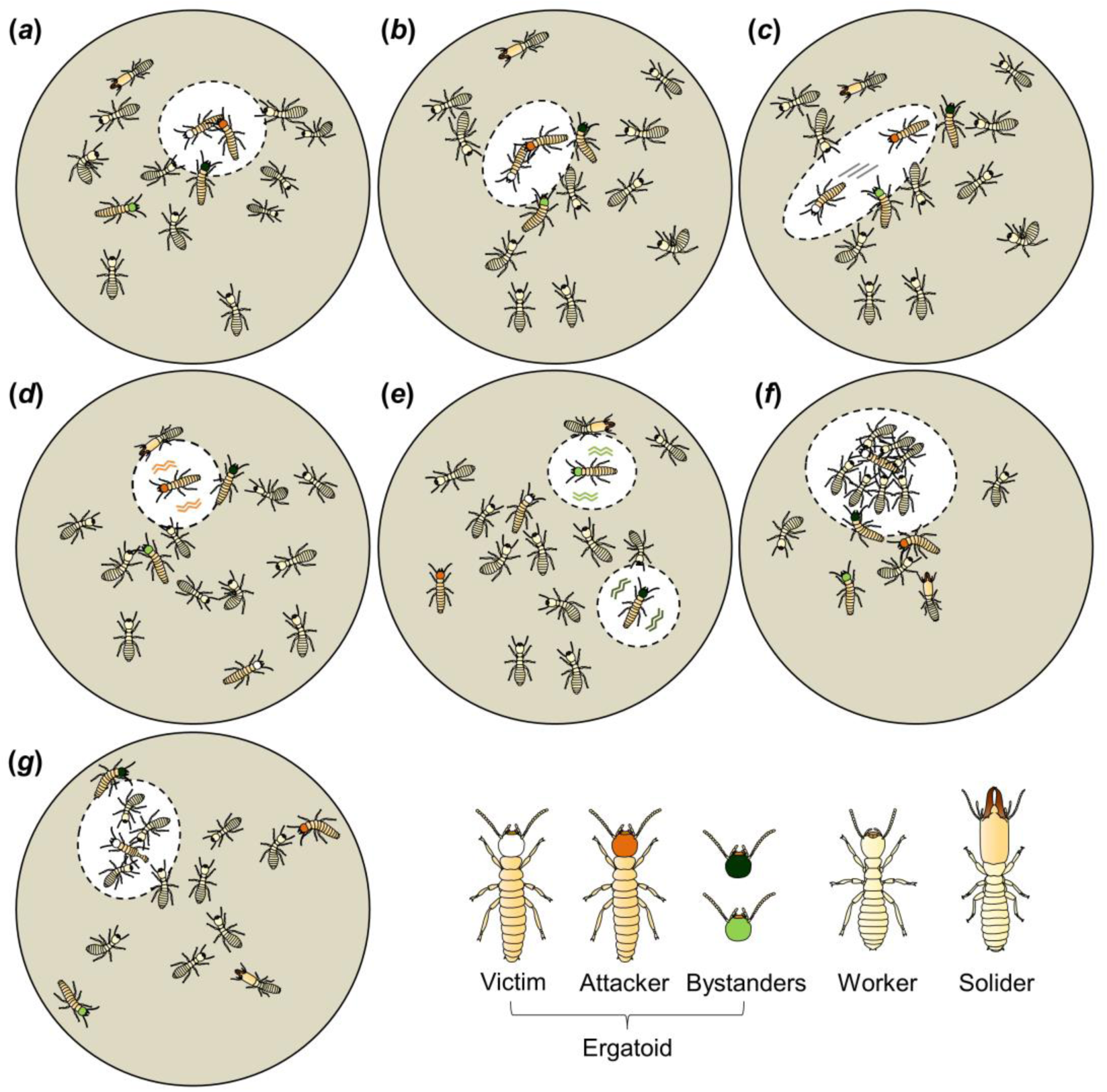
Behavioral process of cooperative policing. (a) An ergatoid attacker holds the victim using mouthparts (Time = −10 sec). (b) The attacker performs a major attack by biting the victim on its abdomen (Time = 0). (c) The major attack causes the victim to quickly flee (Time = 0). (d) The attacker displays alarm behavior by vigorously vibrating its body toward multiple directions (Time = 1 min). (e) Bystanders also display alarm behavior in the same manner (Time = 5 min). (f) Workers then surround and nibble the victim (Time = 30 min). (g) Victim is partially cannibalized (Time = 2 hour). Attacker: ergatoid who performed the first major attack. Victim: ergatoid who received the attack and was later cannibalized. Bystander: other ergatoids.

**Figure 3.**
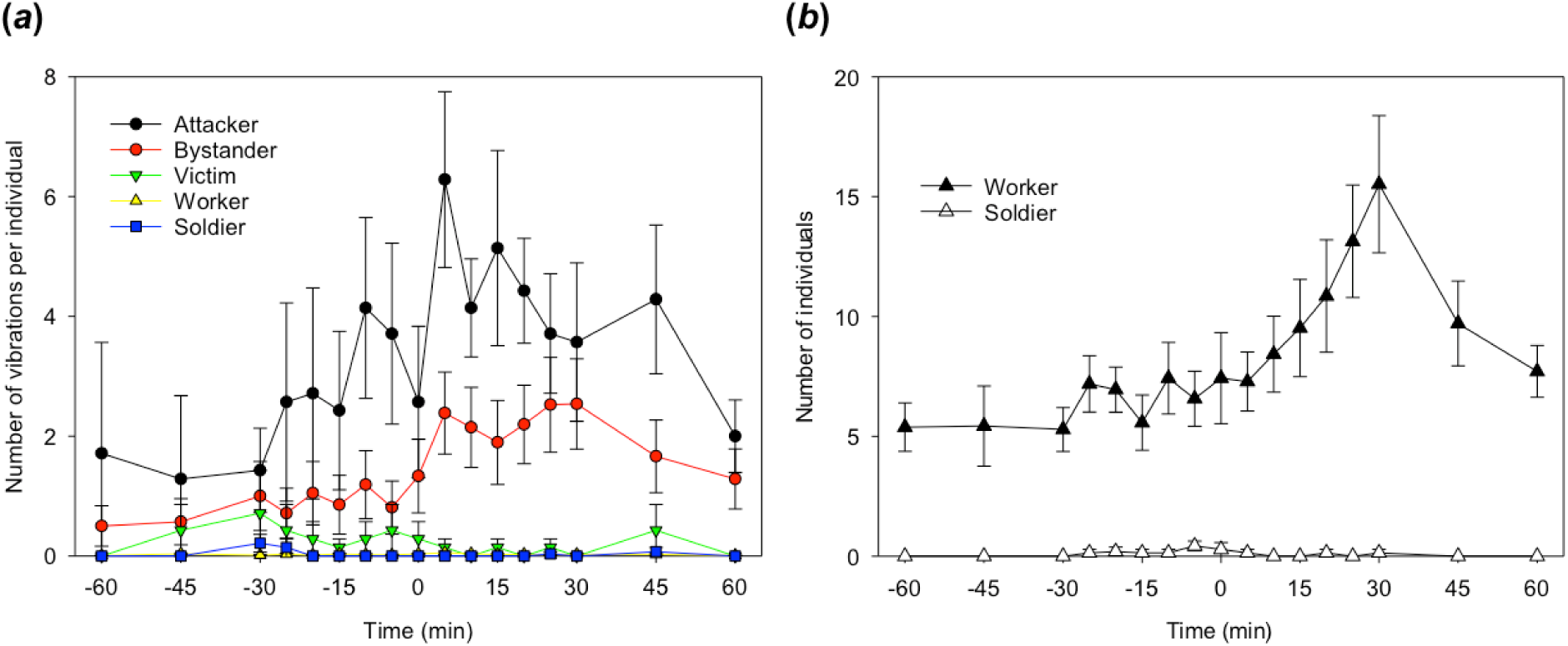
Participation of different castes in policing behavior. (a) Figure shows the number of vibrations per minute performed by each individual (mean ± SE, n = 7). Time “0” was the time of the first major attack. A major attack was recognized when an ergatoid visibly injured another ergatoid. A one-minute sample (30 seconds before and after) was analyzed for each time point. (b) The number of individuals surrounding the victim before and after the attack (mean ± SE, n = 7). The increased density of workers reflects cannibalism.

### (c) Ergatoid elimination is age-dependent

The majority of the aggressive interactions involved an ergatoid attacking the same-sex ergatoid (15 out of 19), while the others involved either a male ergatoid attacking a female (1 out 19), or a group of workers attacking and consuming an ergatoid (1 and 2 cases in female and male treatments, respectively). In the same-sex aggressive interactions, the attacker was always older than the victim in both female and male treatments (Figure 4; female: *Z* = −2.3664, *P* < 0.01, n = 7; male: *Z* = −2.5205, *P* < 0.01, n = 8; Wilcoxon signed-rank test, one-tailed). The median age differences between the pair of attacker and victim were 5 and 2 days for females and males, respectively.

**Figure 4.**
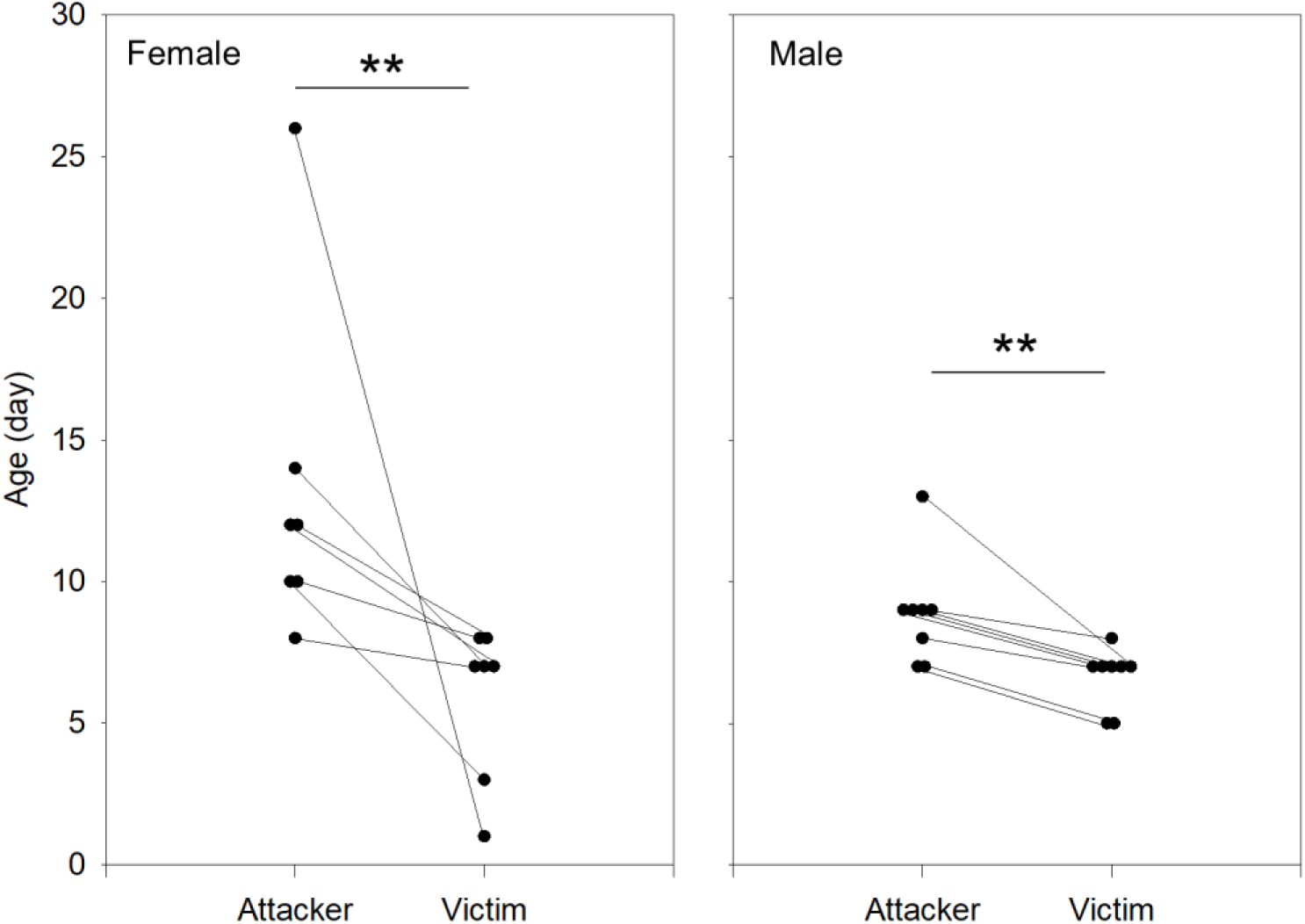
Age of interacting ergatoids in policing behavior. Each dot represents an individual attacker or victim, and lines connect the attacker to the corresponding victim. The negative slope of the lines in both graphs shows that an older ergatoid attacked a younger ergatoid in every replication, and the age differences are significant (female: Z = −2.3664, *P* < 0.01, n = 7; male: Z = −2.5205, P < 0.01, n = 8; Wilcoxon signed-rank test, one-tailed; **, *P* < 0.01).

## 4. Discussion

### (a) Cooperative effort and justification of “policing behavior”

Overall, this study reveals a behavioral mechanism regulating reproductive division of labor during reproductive succession in termites. Unlike butting behavior that is an indicator of reproductive dominance reported in a drywood termite *Cryptotermes secundus* and a dampwood termite *Zootermopsis nevadensis* [37, 38], this behavior directly acts to eliminate reproductive individuals, thus fulfilling the definition of policing [10].

Policing in *R. flavipes* is a sequential behavior cooperatively performed by the older reproductives in concert with workers. This is comparable to the sting smearing behavior in a queenless ant, *Dinoponera quadriceps*, a textbook example of policing behavior [39]. In this ant, the alpha female chemically marks a low-ranking challenger using stinger, causing workers to immobilize the marked individual [40]. Such a cooperative effort allows the alpha to inflict punishment indirectly and maintain her dominant status without fighting. Similarly, in *R. flavipes*, the ergatoid attacker does not kill its rival directly; rather, it induces hemolymph exposure of the victim by biting, and proceeds with an alarm behavior. Workers eventually eliminate the injured individual. The workers are likely recruited by the vibrational signals, and the cannibalistic behavior is potentially caused by the chemical cues in the hemolymph, which remains to be investigated.

### (b) Caste fate conflict

In the absence of reproductives and their inhibitory pheromones, the excess ergatoid production in *R. flavipes* is an expression of caste fate conflict, which also occurs in social Hymenoptera. For example, in the *Melipona* stingless bees, caste is self-determined, and immature females selfishly develop into queens to maximize direct reproduction [41]. Such excess queen production causes depletion of workforce and poses a cost to the colony, leading to a situation known as “tragedy of the commons” [42]. The policing behavior in *R. flavipes* provides an effective solution that prevents unsustainable reproduction upon the loss of former reproductives, and supports colony efficiency. Uncontrolled reproduction causes increased resource demand by the reproductive individuals and their brood, and a balanced ratio of reproductives and workers is often optimal to the colony. For example, in a parthenogenic ant *Platythyrea punctate*, worker to brood ratio is limited, and colonies are incapable of rearing brood produced by additional reproductives [23]. Similarly, an optimal allocation to reproduction is required in the Japanese subterranean termite *R. speratus*, and increased number of queens does not lead to additional reproductive output [43].

In social insects, policing often inhibits reproduction of the focal individual without killing it. In Hymenoptera, policing behavior often enforces workers to stop reproducing and cooperate in brood care. In subterranean termites, however, worker-reproductive differentiation is an irreversible process achieved through at least one molt [25]. Ergatoid reproductives are a non-foraging caste that depends on workers to provide food [44], and the presence of excess ergatoids and the subsequent brood are costly for the colony. Elimination of additional ergatoids avoids future colony investment; cannibalism, in addition, allows the colony to recycle nutrients from the policed individuals and partially rescue the cost that has already occurred upon their differentiation.

### (c) Age-dependent elimination

The age of ergatoids plays an important role in the policing behavior of *R. flavipes*. The result is consistent with a dampwood termite *Porotermes adamsoni*, in which neotenic reproductives that develop earlier have higher survivorship than that differentiate later [45]. Age is also positively correlated with dominance rank in the naked mole-rat *Heterocephalus glaber* [46], in which younger individuals receive more aggression than older ones [47]. In termites, older ergatoids have the first chance to utilize resources from the colony, such as food provided by workers and mating opportunities with existing reproductives. These factors may contribute to the maturity of ergatoids in terms of gonad development [48], body weight, and mandible sclerotization, thus allowing them to outcompete the younger ones. In *C. secundus*, the surviving neotenics performed increased interactions with workers through proctodeal trophallaxis after their differentiation, while the ones eventually eliminated did not [30]. This behavior is possibly associated with age, but it is yet to be tested if neotenics increase trophallaxis as they become older.

The elimination of younger ergatoids in termites is also similar to the selective elimination of small queens in a stingless bee *Schwarziana quadripunctata*, where large fecund queens are favored and dwarf queens tend to be killed by workers [49]. With a policing behavior that strongly acts against small queens, small females should be less tempted to develop into queens, a situation indicated by a theoretical study based on inclusive fitness theory [49]. In *R. flavipes*, while a few ergatoids are formed and eliminated, the majority of workers do not molt into ergatoids. This suggests that the same theory may apply to termites: in the presence of older ergatoid, rather than developing into reproductives and being killed, workers can gain indirect fitness benefits by not differentiating.

### (d) Reproductive competition

The aggressive interaction between ergatoids in termites reflects competition among colony members in inheriting the breeding position after the death of primary reproductives. This form of cooperative policing might be common in termite species where workers or nymphs retain reproductive potential and compete for breeding positions. Indeed, a similar behavior has been observed in a drywood termite, *Kalotermes flavicollis*, where neotenic reproductives attack each other and injured individuals are cannibalized by workers and nymphs [34]. In addition, reproductive competition also occurs upon fusion of neighboring conspecific colonies. In *Z. nevadensis*, reproductives of encountering colonies engage in agonistic behavior, leading to a reduction in their numbers [50]. Moreover, reproductive conflict among unrelated queens happens in species where colonies are founded by pleometrosis. In a fungus-growing termite, *Macrotermes michaelseni*, the mutilation of queen antennae indicates aggression between primary reproductives that are co-founders, and this behavior may influence queen number [51]. These findings suggest that aggressive interaction between reproductives is widespread in termites under diverse social contexts. Aggression represents a conserved component of policing behavior, which, at colony level, regulates reproductive division of labor and serve a collective interest.

### (e) Future directions

Several open questions remain to be addressed in regard to the pheromonal mechanisms underlying policing behavior, including the dynamic change of reproductive pheromones with age, and the chemical cues inducing cannibalism of ergatoids by workers. Importantly, policing behavior, in a broad sense, may be more widespread in termites than previously considered. In addition to a number of convergent traits in social Hymenoptera and Blattodea, such as suicidal colony defense [52], collective foraging using trail pheromones [53], and undertaking behavior [54], policing behavior represents another important behavior that independently evolved in both eusocial groups. Comparative studies on proximate and ultimate aspects of policing behavior among social insect taxa will provide important insights into the evolution of eusociality in insects.

## Supporting information

Raw data

## Supplementary material

Supplementary material includes 1 table, 1 figure, and 4 movies.

## Data accessibility

Raw data can be accessed as supplementary material.

## Authors’ contributions

Q.S., J.D.H. contributed equally to this study. Q.S., J.D.H., K.F.H. and X.Z. designed the study. Q.S. and J.D.H. performed the experiments. Q.S., J.D.H and A.M analyzed the data. Q.S., J.D.H., K.F.H. and X.Z. wrote the manuscript. All authors read and approved the final manuscript.

## Acknowledgments

We thank members of Zhou laboratory for discussion and comments on the study. This work was supported by William L. and Ruth D. Nutting Student Research Grant from the International Union for the Study of Social Insects (North American Section) to Q.S., and a Hatch fund (Accession Number: 1004654; Project Number: KY008071) from the USDA National Institute of Food and Agriculture to X.Z.

