## Supplementary material for "Cooperative policing behavior regulates reproductive division of labor in a termite"

---

---

**Q.S. and J.D.H. contributed equally to this work.**

**[*]Corresponding Author:**

Dr. Xuguo "Joe" Zhou

Department of Entomology

University of Kentucky

S-225 Agricultural Science Center North

Lexington, KY 40546-0091

---

**Short Title:** Policing behavior in termites

**Supplementary Material Information:** 1 table, 1 figure, and 4 movies

**Table S1.** Detailed information for replications analyzed for policing behavior. Among 33 groups of termites recorded, we observed a total of 12 groups showing ergatoids being cannibalized. In 7 groups, the policed ergatoid (victim) was aggressively attacked by another ergatoid (attacker) before workers cannibalized it, with 4 cases being same-sex attacks, 2 cases involving attacking the opposite-sex, and 1 case where sex-identity of victim was undetermined. These 7 cases were used for behavioral analysis. In the other 5 groups, we did not observe ergatoid attacks from the videos, which potentially occurred shortly before the 24-hour video clips that were captured (see Experimental Procedures below for details); we found that the victim was either bitten by an individual worker (in 2 groups) or multiple workers (in 3 groups) before it was cannibalized. The table listed the information of each replication at the event of policing behavior.

| Rep. | Days Orphaned | Caste Composition (Number of Individuals) | | | | | Sex | |
|---|---|---|---|---|---|---|---|---|
| | | Female Ergatoid | Male Ergatoid | Soldier | Pre-soldier | Worker | Attacker | Victim |
| 1 | 75 | 3 | 1 | 1 | 0 | 73 | F | F |
| 2 | 88 | 2 | 1 | 2 | 1 | 79 | M | F |
| 3 | 89 | 2 | 3 | 2 | 0 | 77 | M | M |
| 4 | 84 | 3 | 1 | 1 | 0 | 59 | F | NA |
| 5 | 68 | 4 | 1 | 2 | 1 | 66 | F | F |
| 6 | 77 | 6 | 1 | 4 | 0 | 74 | F | F |
| 7 | 78 | 5 | 1 | 4 | 1 | 74 | M | F |

F: Female.

M: Male.

NA: The ergatoid was not checked for sex identity because it differentiated and policed over night between the routine daily checks.

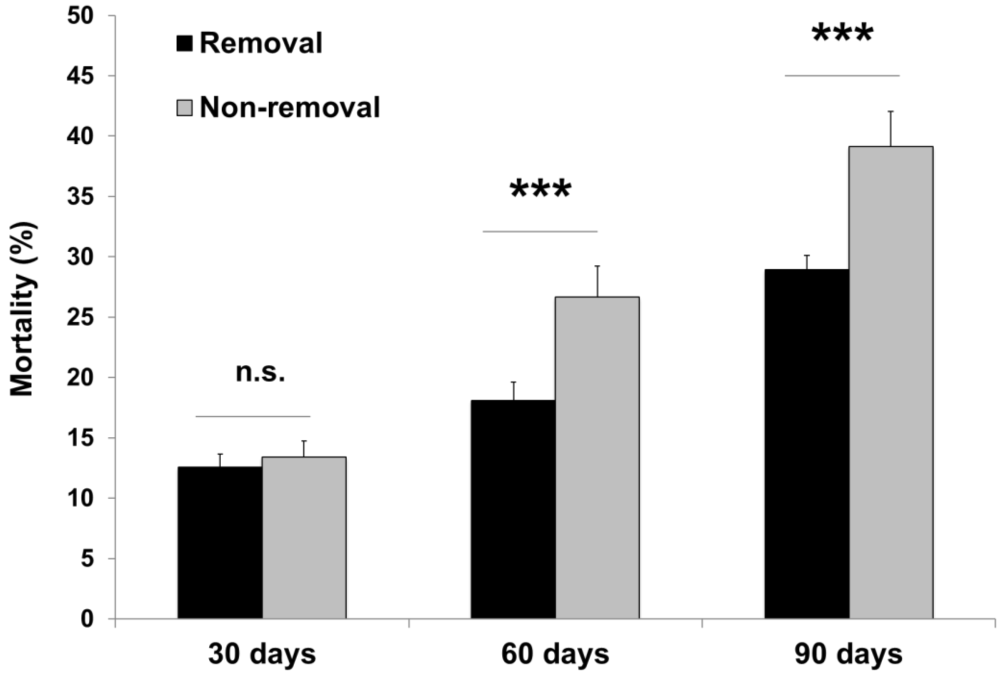


**Figure S1.** Mortalities (%, mean ± SE) post orphaning are shown for removal and non-removal treatments. n.s., not significant; ***, $P < 0.001$; GLMM, Poisson family; n = 20 per treatment per observation day.

**Movie S1.** An ergatoid attacking another ergatoid. The attacker (marked in orange) held the abdomen of the victim (marked in white) with her mouthparts, and took a bite. The victim was injured and quickly fled. Both attacker and victim were female in this clip.

**Movie S2.** Alarm behavior by attacker. After attacking and injuring the victim, the attacker (orange) displayed alarm behavior through vigorous vibrations toward multiple directions.

**Movie S3.** Alarm behavior by bystander. After the event of attack, an ergatoid bystander (dark green) also performed alarm behavior, and examined the injured victim (white) through antennation.

**Movie S4.** Cannibalism by workers. A group of workers were surrounding the victim (white) and consuming it, while the attacker (orange) and a bystander (dark green) were performing alarm behavior.
